# Structural basis for antibody resistance to SARS-CoV-2 omicron variant

**DOI:** 10.1101/2021.12.21.473620

**Authors:** Gabriele Cerutti, Yicheng Guo, Lihong Liu, Zhening Zhang, Liyuan Liu, Yang Luo, Yiming Huang, Harris H. Wang, David D. Ho, Zizhang Sheng, Lawrence Shapiro

## Abstract

The recently reported B.1.1.529 Omicron variant of SARS-CoV-2 includes 34 mutations in the spike protein relative to the Wuhan strain that initiated the COVID-19 pandemic, including 15 mutations in the receptor binding domain (RBD). Functional studies have shown omicron to substantially escape the activity of many SARS-CoV-2-neutralizing antibodies. Here we report a 3.1 Å resolution cryo-electron microscopy (cryo-EM) structure of the Omicron spike protein ectodomain. The structure depicts a spike that is exclusively in the 1-RBD-up conformation with increased mobility and inter-protomer asymmetry. Many mutations cause steric clashes and/or altered interactions at antibody binding surfaces, whereas others mediate changes of the spike structure in local regions to interfere with antibody recognition. Overall, the structure of the omicron spike reveals how mutations alter its conformation and explains its extraordinary ability to evade neutralizing antibodies.

**Highlights:**

- SARS-CoV-2 omicron spike exclusively adopts 1-RBD-up conformation
- Omicron substitutions alter conformation and mobility of RBD
- A subset of omicron mutations change the local conformation of spike
- The structure reveals the basis of antibody neutralization escape

## Introduction

Severe acute respiratory syndrome coronavirus 2 (SARS-CoV-2) emerged as a human pathogen in 2019 in Wuhan, China, causing a disease now known as coronavirus disease 19 (COVID-19) that is characterized by fever, acute respiratory illness, and pneumonia (Callaway et al., 2020; Cucinotta and Vanelli, 2020; Zhou et al., 2020). At the time of writing this article, more than 274 million infections had been reported worldwide, with over 5 million deaths (Dong et al., 2020). Numerous variants have been discovered through sequencing over the past two years, with some major lineages designated as variants of concern (VOCs) due to increased transmissibility, disease severity, resistance to neutralizing antibodies elicited by vaccines, or reduced efficacy of treatments (Planas et al., 2021; Washington et al., 2021). These VOCs are designated alpha, beta, delta and gamma by the World Health Organization, each of which contains a characteristic set of mutations. The Omicron (B.1.1.529) VOC, first detected in southern Africa in November 2021 has spread rapidly to over 60 countries. The astonishingly high transmission rate (R0 >3) and short doubling time (2-3 days) of Omicron cases suggests it could soon become dominant (Burki, 2021). The alarming number of mutations in the spike protein (34), including at least 15 in the receptor-binding domain (RBD), the primary target for neutralizing antibodies, results in substantially compromised efficacy of vaccines and therapeutic antibodies (Liu et al., 2021). Elucidating the structural basis of viral escape becomes a high priority for understanding viral evolutionary pathways and developing new therapeutics.

SARS-CoV-2 utilizes a highly glycosylated spike protein (S) to mediate entry into host cells. S, a class I fusion protein, forms a trimer that adopts a metastable prefusion conformation that undergoes large structural rearrangements in fusion of the host and viral cell membranes (Bosch et al., 2003; Shang et al., 2020). Host cell ACE2 receptor binding is thought to destabilize the prefusion trimer, leading to shedding of the S1 subunit and transition of S2 to an elongated helical postfusion conformation (Benton et al., 2020; Cai et al., 2020; Hoffmann et al., 2020; Wang et al., 2020). The RBD of S1 undergoes conformational motions between an “up” state where the ACE2 receptor binding site is accessible, and a “down” state where it is hidden (Walls et al., 2020; Wrapp et al., 2020).

Neutralizing antibodies most often target the domains at the top of the spike – RBD (Barnes et al., 2020; Brouwer et al., 2020; Cao et al., 2020; Cerutti et al., 2021c; Chen et al., 2020; Ju et al., 2020; Liu et al., 2020b; Pinto et al., 2020; Rapp et al., 2021; Shi et al., 2020; Tortorici et al., 2020; Zost et al., 2020), and NTD (Cerutti et al., 2021a; Cerutti et al., 2021b; Chi et al., 2020; McCallum et al., 2021a; Suryadevara et al., 2021). The large number of mutations in Omicron has raised questions about its evolutionary origin. Some have proposed that it developed in a chronically infected individual with an impaired immune response, whereas others think it developed in parallel with other variants in a population not monitored by sequencing (Kupferschmidt, 2021). Mutations in VOCs accumulate mainly in the S protein. Omicron has 15, 8, and 11 mutations in RBD, NTD, and S2 subunits respectively, with 6 of the 34 mutations observed in other VOCs or variants of interest. The RBD mutations include G339D, S371L, S373P, S375F, K417N, N440K, G446S, S477N, T478K, E484A, Q493R, G496S, Q498R, N501Y, and Y505H. Some of these mutations have known functional consequences, such as K417N, S447N, E484A, Q493R, which contribute to immune escape (Harvey et al., 2021; Wang et al., 2021b), and N501Y which contributes to higher infectivity (Tian et al., 2021). But other Omicron mutations in RBD and other domains have unknown functional impact, individually or in combination.

Here we present a cryo-EM structure of the Omicron spike, which adopts an exclusively 1-up RBD conformation. The overall architecture of the spike is conserved, but the mobility of RBDs appears increased over other variants. Surface differences appear to be localized around sites of antibody recognition, with serious implications for immune evasion.

## Results

### Cryo-EM structure of omicron spike

For structure determination, we produced a soluble version of the Omicron spike corresponding to residues 1 to 1208 of the ectodomain, and including two proline mutations in S2 which have been previously used to stabilize the spike in its prefusion form (Walls et al., 2020; Wrapp et al., 2020), and a C-terminal His tag. The protein was expressed in HEK-293 cells and purified by His-tag affinity chromatography, and this protein was used for the preparation of cryo-EM grids. The spike appeared to show a slight preferred orientation on the cryo-EM grids, so we collected data with a 30° tilt angle. We collected and processed 13695 cryo-EM movies to obtain a 3D reconstruction at 3.1 Å resolution (**Figure 1A, Figure S1, Table S1**).

**Figure 1.**
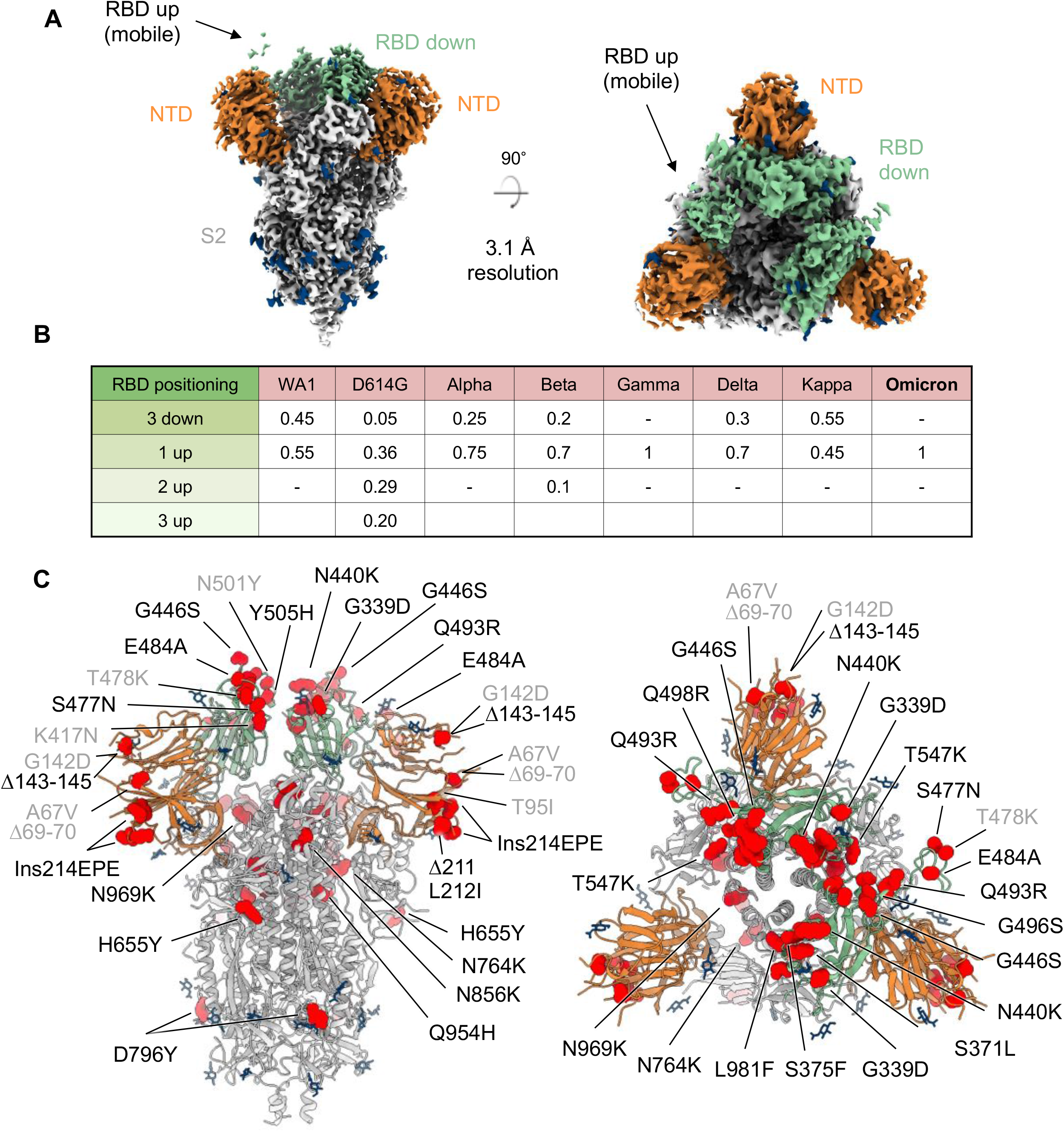
Cryo-EM structure of prefusion SARS-CoV-2 omicron (B.1.1.529) spike. (A) Cryo-EM map of SARS-CoV-2 omicron S2P spike in the prefusion state shown in two orthogonal views. The density for the single RBD up is barely visible at the optimal contour level due to high mobility of the domain. NTD is colored in orange, RBD in green, glycans in blue, the rest of the trimer in gray. (B) Relative population of RBD states observed in cryo-EM structures of SARS-CoV-2 spike for different variants. (C) Structure of SARS-CoV-2 omicron spike in the 1 RBD up state with mutations highlighted in red. Mutations observed in previous variants are labelled in gray, new omicron mutations are labelled in black. See also Figure S1, Figure S2, Table S1.

The overall structure is similar to the Wuhan spike, with the totality of particles in the 1-RBD up conformation. The absence of electron density for the RBD in the up conformation is a result of its high mobility, as confirmed by 3D variability analysis. The main variability component observed within the final particle set showed an oscillatory motion for the RBD up, coupled to breathing of the two RBDs in the down position (**Figure S2**). The electron densities for the two RBDs down were not equivalent, with the best RBD density observed for the down-RBD located between the other down-RBD and the up-RBD.

The single 1-RBD-up conformation observed for omicron is also typical of the gamma variant (Wang et al., 2021a; Zhang et al., 2021b), while for other variants an equilibrium of different states has been reported (**Figure 1B**) (Gobeil et al., 2021; Yurkovetskiy et al., 2020; Zhang et al., 2021a; Zhang et al., 2021b). Specifically, the disappearance of the RBD up density observed in omicron spike was reported for the alpha variant (Gobeil et al., 2021).

Most of the omicron mutations were visible in the cryo-EM structure and their location in the context of spike is depicted in **Figure 1C**. Mutations Δ69-70 (NTD), S373P (RBD), N679K and P681H (proximal to the S1/S2 cleavage site) belonged to flexible regions that could not be resolved in the cryo-EM structure. The remaining 30 mutations were clearly visible in the cryo-EM structure. The RBD mutations are mostly clustered at the inter-protomer RBD-RBD interface, while the NTD mutations are located in the flexible loops distal from the trimer axis. The S2 mutations are mostly located at the top of the subunit, at the interface with S1.

### Similarity and difference between omicron and D614G spike

To evaluate whether the omicron mutations induce overall orientation changes among spike domains, we superimposed the structures of omicron variant to the D614G wildtype with 1-up RBD (PDB ID 7KRR) (**Figure 2A**). The comparison revealed an overall root mean square deviation (RMSD) of 1.1 Å and 0.6 Å for S1 and S2 subunits respectively. The measured distance between NTD domains of the three protomers showed that the NTD domain from protomer A (NTD_A_), which has an up-RBD, is 5 Å closer to the NTD domain of protomer B (NTD_B_) than that of the D614G spike (**Figure 2B**). We also observed that the S2 helix bundle (residues 988 to 1033) packed tighter between protomers A and C than the wildtype spike (**Figure 2C**). This conformation change may be associated with two mutations: T547K and L981F, both of which introduce larger side chains that could increase the repulsion between the helix bundle and adjacent domains.

**Figure 2.**
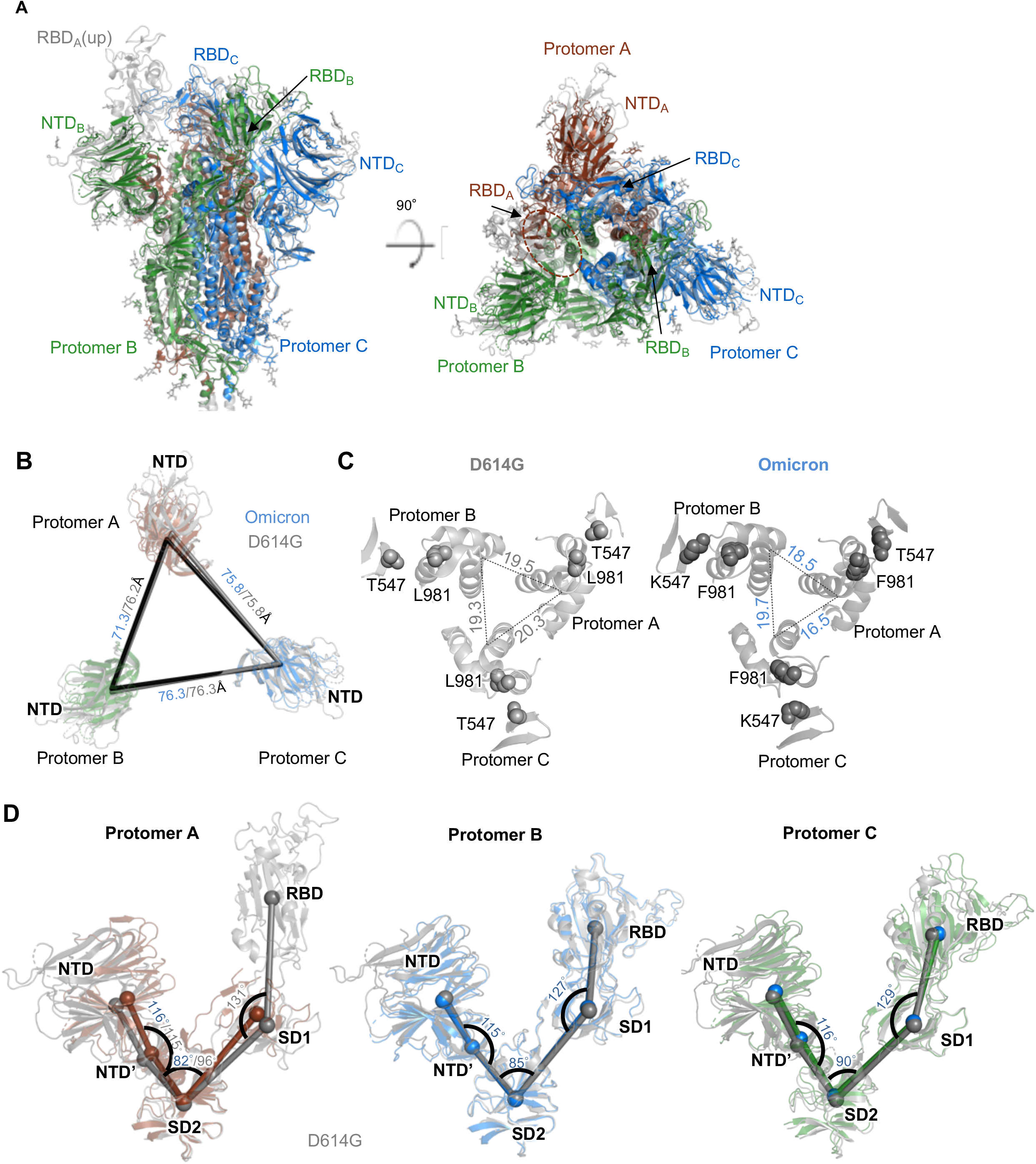
Structural comparison of SARS-CoV-2 omicron spike with D614G wildtype. (A) Superposition of omicron spike with D614G spike. The S2 subunit is used for superimposition. (B) Distance between NTD domains of omicron spike and D614G spike. (C) The inter-protomer distance between S2 helices in omicron is shorter than what observed in D614G spike. (D) Measured angles between NTD, NTD’, SD2, SD1, and RBD showed that protomer A with an up-RBD has altered angles between NTD’, SD2, and SD1. The two protomers with RBDs down show similar domain orientation. Thus, only angles of the omicron spike are shown.

We then determined the center of mass (COM) for NTD, NTD’, SD2, SD1, and RBD and used COMs to calculate angles between these domains. The result revealed that protomer A has a smaller angle between NTD’, SD2, and SD1, while the angles between other domains are highly similar to the wildtype spike (**Figure 2D**). The angles between the five domains in protomer B and C have no difference compared to the wildtype. We then measured the buried accessible surface area (bASA) between the above domains and found that almost every domain-domain interface bASA increases slightly in omicron compared to the wildtype (**Table S2**). Remarkably, NTD has a 3-fold increased bASA with adjacent RBD, coupled to a 2 Å reduced distance between RBD and NTD through the orientation change in NTD. In summary, our analyses showed tighter packing of the omicron spike with increased asymmetry.

### Effects of omicron mutations on spike conformation

We next mapped the omicron mutations to the spike structure and assessed their potential effects on spike conformation. The majority of the RBD mutations are located in the receptor binding motif (RBM), inner side, and outer side epitope regions (**Figure 3A**). The superimposition of the omicron and wildtype RBDs showed an RMSD of 0.75 Å. We then calculated Cα distance for each RBD residue between omicron and wildtype, and observed that six mutations (S371L, S373P, S375F, G446S, S477N, and T478K) are located at regions with Cα distance larger than 2 Å (**Figure 3B**), suggesting that these mutations may account for the conformational change in the omicron RBD. In particular, we observed that mutations S371L, S373P, and S375F not only alter the conformation of loop 371-376 but also result in the motion of helix 365-370 closer to helix 337-343, which may alter the conformation of the N-linked glycosylation at N343 (**Figure 3C** left panel). The formation of new hydrogen bond networks by mutations G446S, G496S, Q498R, and N501Y stabilize loop 443-451 to a new conformation (**Figure 3C** right panel).

**Figure 3.**
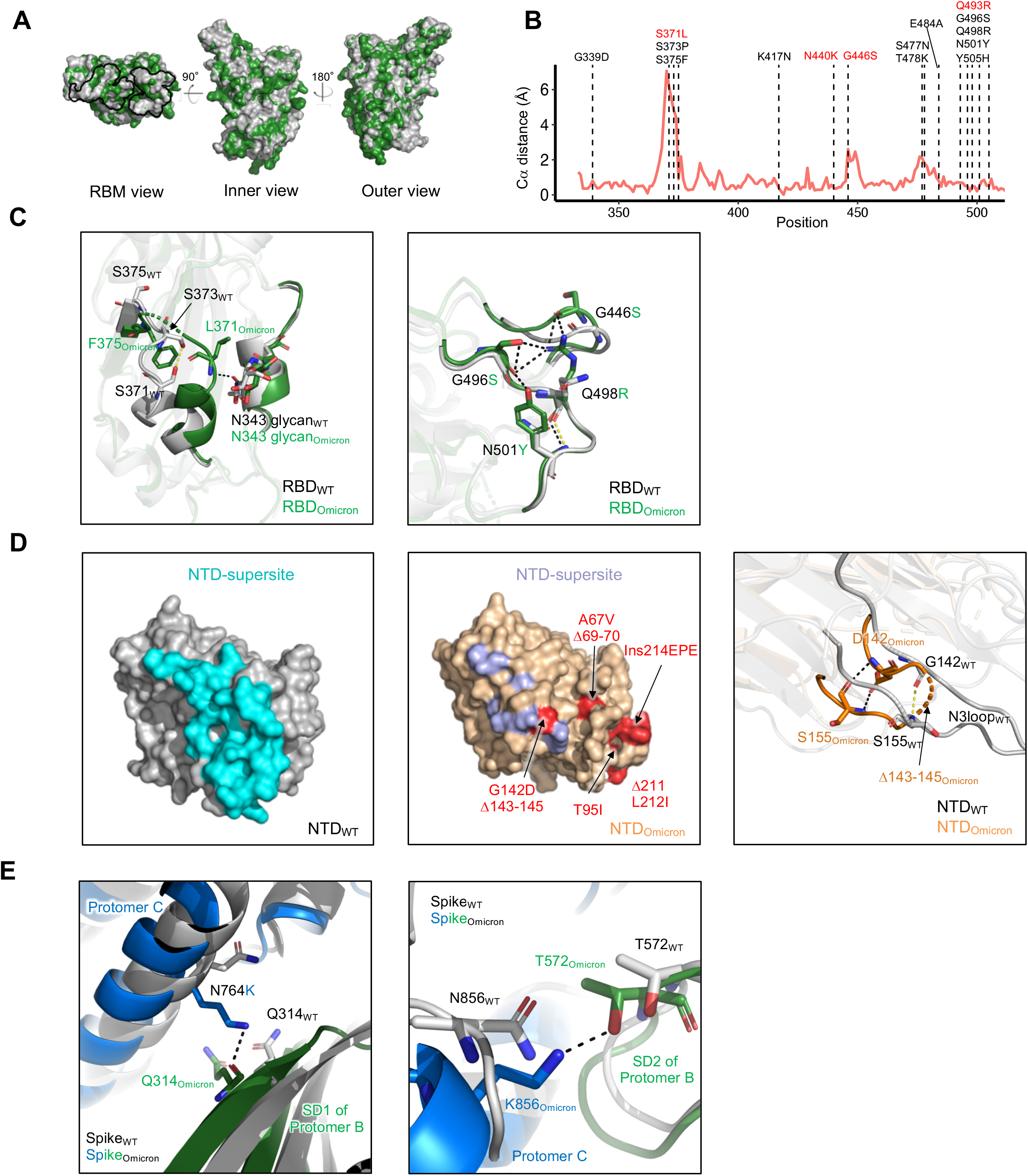
Omicron spike mutations alter local conformation and polar interaction pattern. (A) Superposition of omicron RBD with WT RBD. The Omicron RBD is colored in green and WT RBD in gray. The dark line shows the footprint of the receptor binding motif (RBM). (B) Per residue Cα distance between omicron and WT RBD. RBD mutations in omicron are labeled. The omicron specific mutations labeled in red resulted in dramatic and broad antibody neutralization resistance. (C) Details of conformation changes in omicron RBDs. Left panel, 367-375 conformation change. Right, 444-448 loop conformation change. Black and yellow dashed line show hydrogen bonds in omicron and WT RBDs respectively. (D) Comparison of omicron and WT NTD. Left panel, structure for WT NTD with the NTD supersite colored cyan. Middle panel, structure for omicron NTD, the blue residues show the NTD supersite on omicron, the red residues labeled the omicron mutations. Right panel, details of N3 loop change in omicron compare with WT. (E) S2 mutations N764K and N856K in omicron form new polar interactions with SD2 and SD1 from adjacent protomers respectively. See also Figure S3

The majority of NTD mutations are located at the antigenic supersite targeted by most NTD-directed neutralizing antibodies. Our omicron structure revealed substantial conformational changes in the NTD supersite (**Figure 3D** left and middle panels). We also determined part of the N3 loop with G142D and Δ143-145 (**Figure 3D** right panel). In addition, in the two RBD down protomers, we observed omicron S2 mutations N764K and N856K to form new hydrogen bonds with SD1 and SD2 domains from adjacent protomers respectively (**Figure 3E**). Two conserved residues nearby these mutations also form additional hydrogen bonds in omicron spike (**Figure S3C**). Because both SD1 and SD2 undergo a substantial rearrangement when the RBD switches to the up conformation, these interactions may help stabilizing the RBD in the down conformation by locking SD1 and SD2, an effect similar to other S2 mutations (Gobeil et al., 2021).

### Mechanisms of antibody escape

To understand the structural basis of immune evasion by omicron, we analyzed the effects of mutations substantially contributing to neutralization reduction of major RBD and NTD antibody classes that represent convergent human antibody response to SARS-CoV-2. Consistently with previous studies (Amanat et al., 2021; Wang et al., 2021b), K417N and Q493R impair neutralization of class 1 and 2 antibodies through steric clash and reduced polar interactions (**Figure 4A**, **Figure S4A**). E484A reduces polar interactions with class 2 antibodies. For class 3 antibodies, G446S induces steric clashes with CDRH3. The omicron structure also revealed that the conformational changes in loop 443-451 enhance steric hindrance (**Figure S4B** left panel). For class 4 antibodies, the altered conformation of helix 365-370 and loop 371-376 reduces the interaction with the CDRH3 tip of class 4 antibodies (**Figure 4B**, **Figure S4B** right panel), which may increase the entropy of the epitope/paratope interaction. In addition, the mutations at 371, 373, and 375 are likely to impair recognition of quaternary epitope recognizing class 1 or 2 antibodies (**Figure S4C**). The NTD point mutations and deletions alter the antigenic supersite substantially (**Figure 4C**), abolishing neutralization by most NTD-directed neutralizing antibodies (Liu et al., 2021).

**Figure 4.**
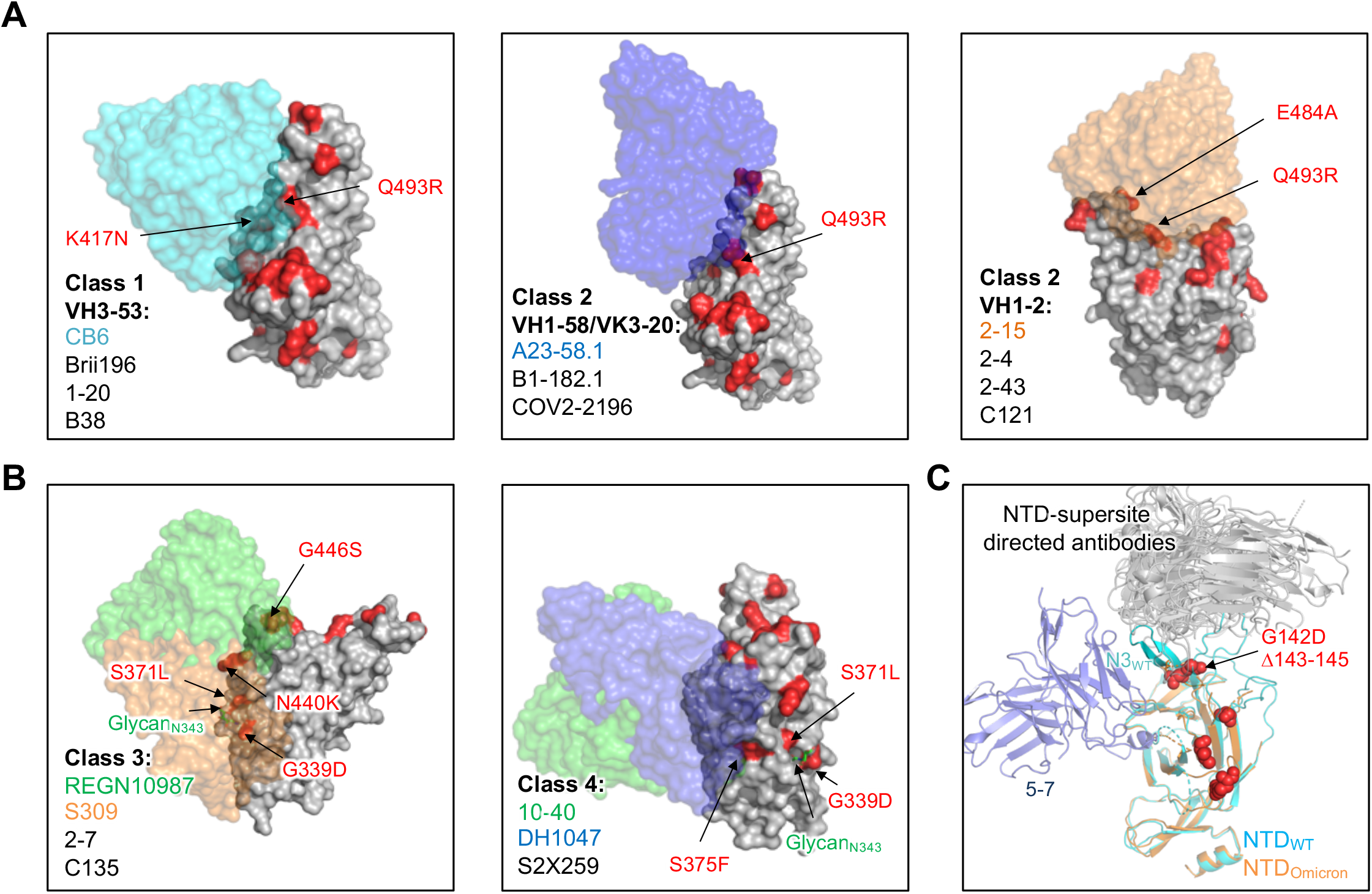
Structural basis of neutralizing antibody escape by omicron. (A) Surface diagram of class 1 and 2 antibodies bound to RBD. Left panel, VH3-53 derived antibodies with CB6 as an example. Middle panel, VH1-58/VK3-20 derived antibodies with A23-58.1 binding mode shown. Right panel, VH1-2 derived antibody with 2-15 binding mode shown. Mutations in omicron RBD are colored in red. (B) Surface diagram of the class 3 and 4 antibodies bound to omicron RBD. (C) Cartoon diagrams of the omicron and WT NTDs in complex with antigenic supersite-directed antibodies and 5-7. Omicron mutations are shown as red spheres. See also Figure S4.

## Discussion

We report the cryo-EM structure of the SARS-CoV-2 spike of the highly transmissible omicron variant, which includes an unprecedented number of mutations and achieves an unprecedented level of antibody escape. While the structure is similar overall to D614G spike, it exists in an all 1-RBD-up conformation with increased mobility of RBD. Many mutations in the omicron spike alter antibody binding surfaces, but some mutations change the local conformation of spike to evade antibody recognition.

The observed 1-RBD-up spike conformation may have evolutionary advantage. For example, the cryptic epitopes, which are only available in the up-RBD and are recognized by antibody classes 1 and 4, have a lower frequency of exposure compared to the wildtype. Together with increased mobility of the up-RBD and mutations, the omicron spike can shed off recognition by dominant human antibodies elicited by infection and vaccination. Since many antibodies recognize two up-RBDs simultaneously to enhance neutralization through avidity, the reduced number of up-RBDs may decrease this effect (Liu et al., 2020a; Rapp et al., 2021). In this study, we observed that S2 mutations N764K and N856K may play a role in stabilizing RBDs in the down conformation through additional interactions with SD1 and SD2 domains. L547K and L981F are close to the RBD in the down conformation of adjacent protomers (**Figure S3D**), which may also contribute by altering S2/RBD interactions.

A convergent antibody response in humans results in distinct spike regions subject to antibody neutralization. Antibodies directed against RBD embody one of the primary immune defenses against SARS-CoV-2. Omicron spike contains 15 mutations in RBD, many overlapping the recognition surfaces of neutralizing antibodies. Compared to other VOCs, omicron also accumulated mutations extensively within and surrounding the receptor binding motif. These mutations lead to the escape of antibodies of all major structural classes - including classes 1, 2 and 3 - which has been observed to a lesser extent with other VOCs. Differently, omicron has two unique clusters of mutations in the RBD. One cluster is G446S, G496S, and Q498R. This cluster of mutations cooperatively alter the loop 443-451 recognized by classes 1, 2, and 3 antibodies, suggesting the result of strong immune selection pressure. The second mutation cluster is S371L, S373P, and S375F, which alter the conformation of loop 371-376 and helix 365-370. These mutations allow omicron to evade class 4 antibodies (Saunders et al., 2021), directed to a highly conserved region on the ‘side’ of RBD, which has so far been observed with no other VOC. Besides, two additional classes of antibodies, quaternary-epitope recognizing class 1 and 2 antibodies (2-43 and S2M11, etc.) (Rapp et al., 2021; Tortorici et al., 2020) and S309-like class 3 antibodies (Andreano et al., 2021) are also impaired by the three mutations. In addition, helix 365-370 tends to have stronger interaction with the base of the N-linked glycosylation at N343, which plays roles in stabilizing two adjacent down-RBDs (Rapp et al., 2021). Together with G339D, these mutations may alter the orientation of the N-linked glycosylation at N343, which may affect antibody binding to adjacent epitope regions.

NTD-directed antibodies also constitute an important defense, with most neutralizers directed to a single antigenic supersite (Cerutti et al., 2021b; McCallum et al., 2021a). Similar to other VOCs, omicron contains mutations and deletions directly within loops of the supersite providing a structural explanation for escape from this major class of antibodies (McCallum et al., 2021b). Overall, this is consistent with the idea that omicron evolved in response to the immune pressure of neutralizing antibodies.

In summary, this study reports the structural impact of mutations emerged with the omicron variant on the architecture of SARS-CoV-2 spike. The cryo-EM structure reveals the details of omicron spike conformational modulation and immune evasion in the context of the natural evolution of the virus. As the omicron mutations overlap with epitopes of neutralizing antibodies, our structural analysis explains the antibody resistance and informs the identification of effective therapeutic and prophylactic strategies.

## Acknowledgements

Cryo-EM data collection was performed at the Columbia University Cryo-Electron Microscopy Center. Support for this work was provided by the fund UR010655/70003/ZS2248 to ZS.

## Author Contributions

G.C. determined the cryo-EM structure of SARS-CoV-2 omicron spike; Y.G. and Z.S. performed bioinformatics analyses; L.L. produced SARS-CoV-2 omicron spike; Z.Z. collected cryo-EM data; D.D.H. supervised spike production; L.S. supervised cryo-EM study; Z.S supervised informatics studies. L.S. and Z.S. oversaw the project and –with G.C. and Y.G. – wrote the manuscript, with all authors providing revisions and comments.

## RESOURCE AVAILABILITY

### Lead Contact

Further information and requests for resources and reagents should be directed to and will be fulfilled by the Lead Contact, Lawrence Shapiro.

### Materials Availability

Expression plasmids generated in this study for expressing SARS-CoV-2 protein will be shared upon request.

### Data and Code Availability

Cryo-EM maps and fitted coordinates of Omicron spike are in the process of being deposited to the EMDB and RCSB PDB respectively.

## EXPERIMENTAL MODEL AND SUBJECT DETAILS

### Cell lines

Expi293 cells were from ThermoFisher Scientific Inc (ThermoFisher, cat#A14527 and cat# A39240 respectively). HEK 293T/17(cat# CRL-11268™) and Vero E6 cells (cat# CRL-1586™) were from ATCC.

## METHOD DETAILS

### Expression and Purification of SARS-CoV-2 Spike

The ectodomain with 2P and furin mutations of SARS-CoV-2 B.1.1.529 trimer was synthesized, fused to an 8 × His tag at the C terminus and then cloned into the paH vector. To purify the S trimer protein, the expression vector was transiently transfected into Expi293 cells using FectoPRO (Polyplus-transfection SA). Two days after transfection, the S trimer protein was purified using Ni-NTA resin (Invitrogen).

### Cryo-EM sample preparation

The sample for cryo-EM analysis of SARS-CoV-2 S2P omicron spike was concentrated to 0.5 mg/mL final trimer concentration. To prevent aggregation during vitrification, 0.005% (w/v) n-dodecyl β-D-maltoside was added to the sample prior to plunge freezing. Cryo-EM grids were prepared by applying 2 μL of sample to a freshly glow-discharged UltrAuFoil gold grid 0.6/1 300 mesh; the sample was vitrified in liquid ethane using a Vitrobot Mark IV with a blot time of 3 s.

### Cryo-EM data collection, processing and structure refinement

Cryo-EM data were collected using the Leginon software (Suloway et al., 2005) installed on a Titan Krios electron microscope operating at 300 kV, equipped with a Gatan K3-BioQuantum direct detection device. The total dose was fractionated for 2.5 s over 50 raw frames. Processing of the first 1000 micrographs showed a slight preferred orientation in the 2D classes. The following micrographs were collected applying a 30° tilt (Tan et al., 2017). Motion correction, CTF estimation, particle picking, extraction, 2D classification, ab initio model generation, 3D classification, 3D refinements and local resolution estimation were carried out in cryoSPARC 3.2 (Punjani et al., 2017) The final 3D reconstruction was obtained using non-uniform refinement with C1 symmetry, achieving a resolution of 3.1 Å. 3D variability analysis (Punjani and Fleet, 2021) was performed to confirm the absence of RBDs down conformations and to sample the mobility of the final particle set in distinct states. 3D classification using the two extremes of the variability mode did not improve the quality of the map, and the particles were merged in a consensus refinement. Particles were symmetry-expanded in C3 to produce a locally refined map using a mask built around the NTD, subsequently used to refine the NTD model and visualize most of its mutations.

The structural model of SARS-CoV-2 spike PDB entry 6VYB (Walls et al., 2020) was used as initial template for model building of the trimer. PDB entries 7EAM (Li et al., 2021) and 7L2C (Cerutti et al., 2021b) were used as initial templates to build the RBD and the NTD respectively. Automated and manual model building were iteratively performed using real space refinement in Phenix (Adams et al., 2010) and Coot (Emsley and Cowtan, 2004) respectively. Half maps were provided to Resolve Cryo-EM tool in Phenix to support manual model building (Terwilliger et al., 2020). Geometry validation and structure quality assessment were performed using EMRinger (Barad et al., 2015) and Molprobity (Davis et al., 2004). Map-fitting cross correlation (Fit-in-Map tool) and figures preparation were carried out using PyMOL and UCSF Chimera (Pettersen et al., 2004) and Chimera X (Pettersen et al., 2021). A summary of the cryo-EM data collection, reconstruction and refinement statistics is shown in Table S1.

The cryo-EM structural model and maps are in the process of being deposited in the RCSB PDB and EMDB.

### Calculation of domain angles and distance and identification of domain interfaces

PyMOL was used to perform the angle and distance calculations and generate plots (Schrodinger, 2015). For each domain, we selected a set of residues and used their Cα for determining the center of mass of the domain. The residues used are NTD (residues 27 to 69, 80 to 130, 168 to 172, 187 to 209, 216 to 242, and 263 to 271), NTD’ (residues 44 to 53 and 272 to 293), RBD (residues 334 to 378, 389 to 443, and 503 to 521), SD1 (residues 323 to 329 and 529 to 590), SD2 (residues 294 to 322, 591 to 620, 641 to 691, and 692 to 696) (Gobeil et al., 2021). PISA was used to identify interface residues, as well as calculate buried accessible surface area and identify polar interactions (Winn et al., 2011).

## QUANTIFICATION AND STATISTICAL ANALYSIS

Cryo-EM data were processed and analyzed using CryoSparc. Cryo-EM structural statistics were analyzed with Phenix and Molprobity. Statistical details of experiments are described in Method Details or figure legends.

---

**Figure S1.**
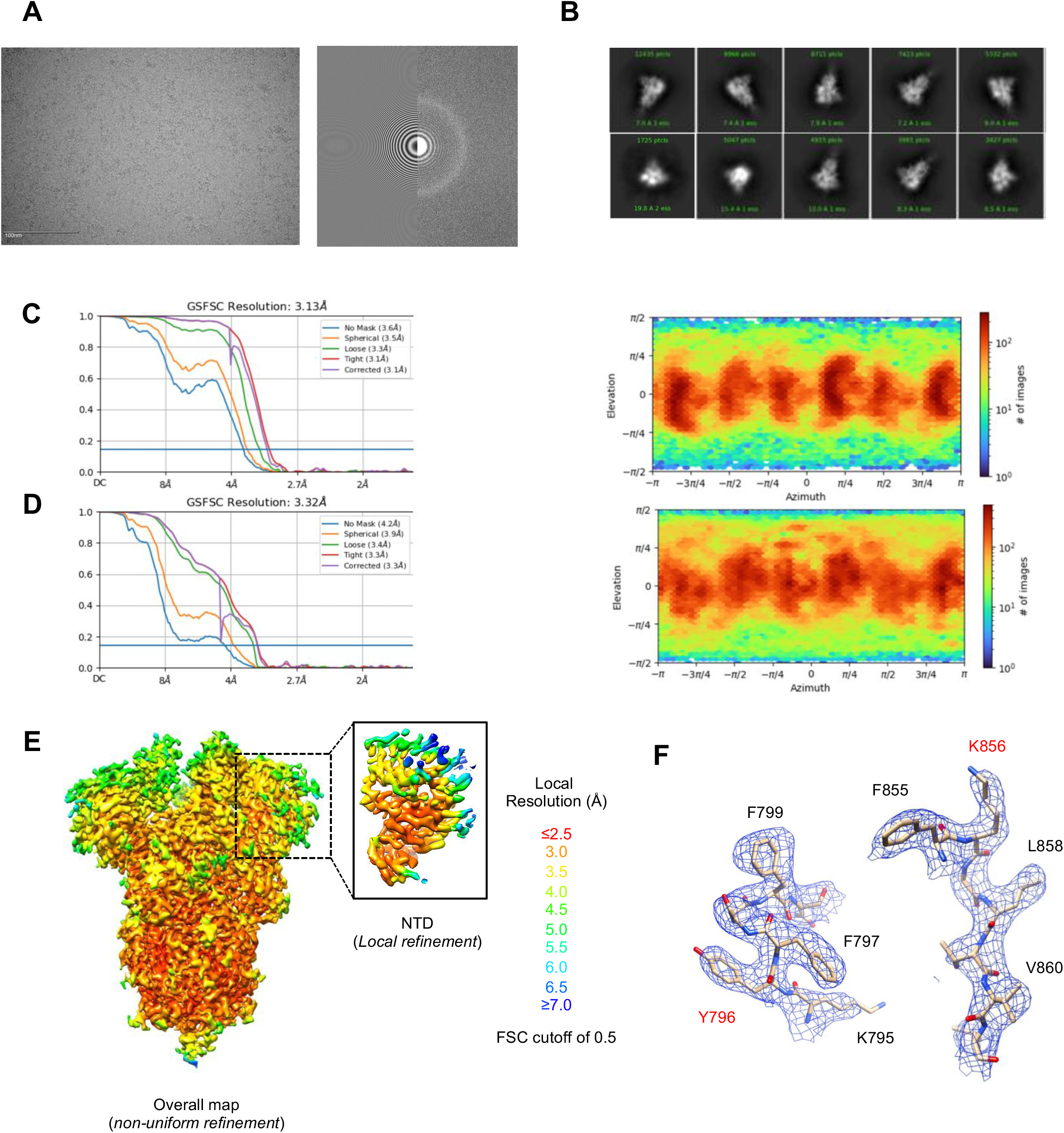
Cryo-EM details of SARS-CoV-2 S2P spike omicron variant, Related to Figure 1. (A) Representative micrograph and CTF of the micrograph are shown. (B) Representative 2D class averages are shown. (C) The gold-standard Fourier shell correlation resulted in a resolution of 3.13 Å for the overall map using non-uniform refinement (left panel). The orientations of all particles used in the final refinement are shown as a heatmap (right panel). (D) The gold-standard Fourier shell correlation resulted in a resolution of 3.32 Å for the local refinement of NTD; particles were symmetry expanded in C3 and aligned using a mask comprising the NTD. The orientations of all particles used in the final refinement are shown as a heatmap (right panel). (E) The local resolution of the final overall map and locally refined map for NTD are shown, generated through cryoSPARC using an FSC cutoff of 0.5. (F) Representative density is shown for the surroundings of mutated residues (red label) in the S2 subunit. Carbon atoms are colored in beige, oxygen in red, nitrogen in blue.

**Figure S2.**
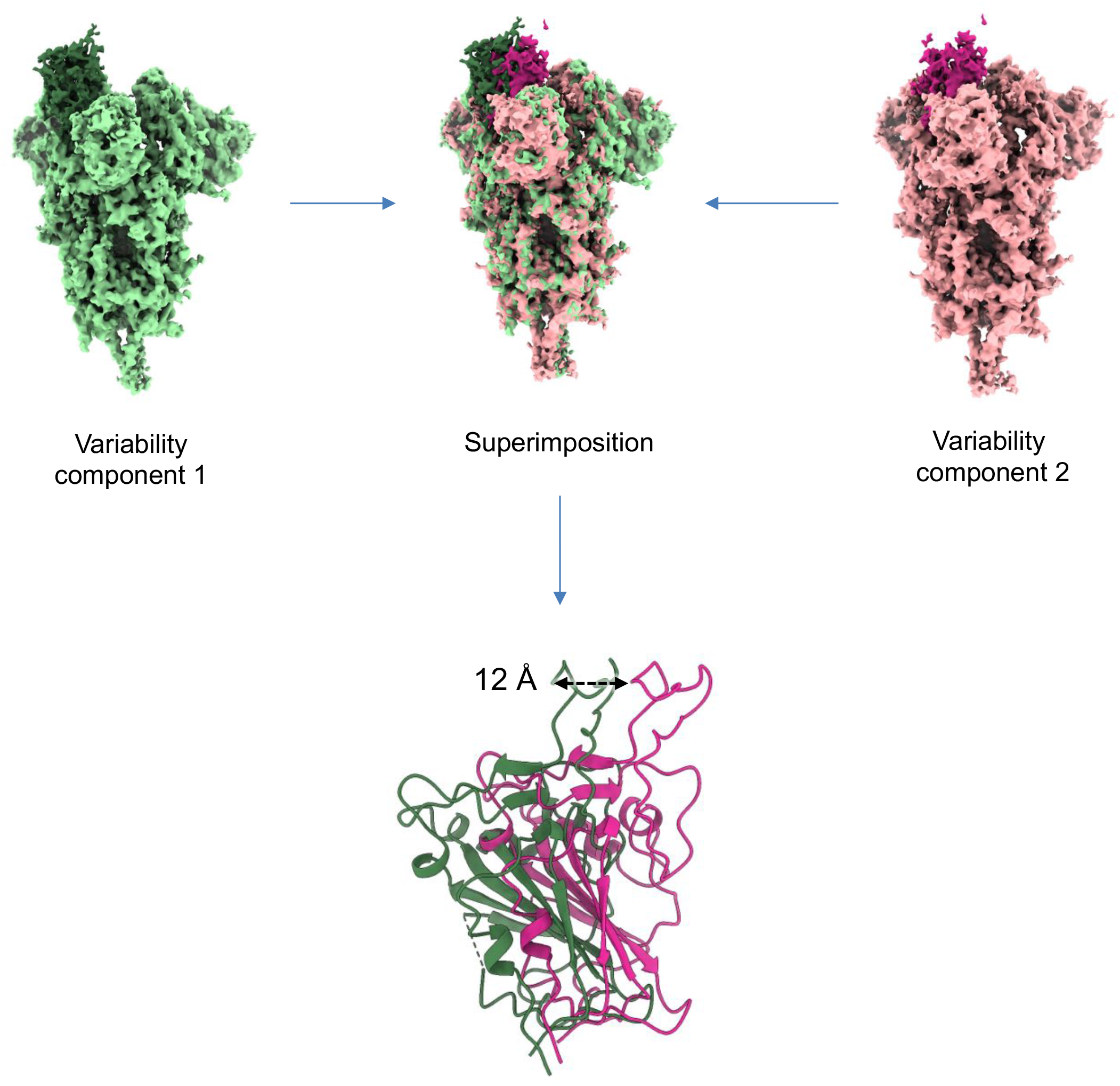
Mobility of RBD in the up conformation, Related to Figure 1. The two maps representing the extremes of the 3D variability observed in the 1 RBD up conformation of SARS-CoV-2 omicron spike are shown in green (RBD dark green) and pink (RBD magenta) together with their superimposition. Fitting of RBD models in the maps shows an oscillatory motion resulting in a maximal 12 Å displacement for the “hook” region of RBD.

**Figure S3.**
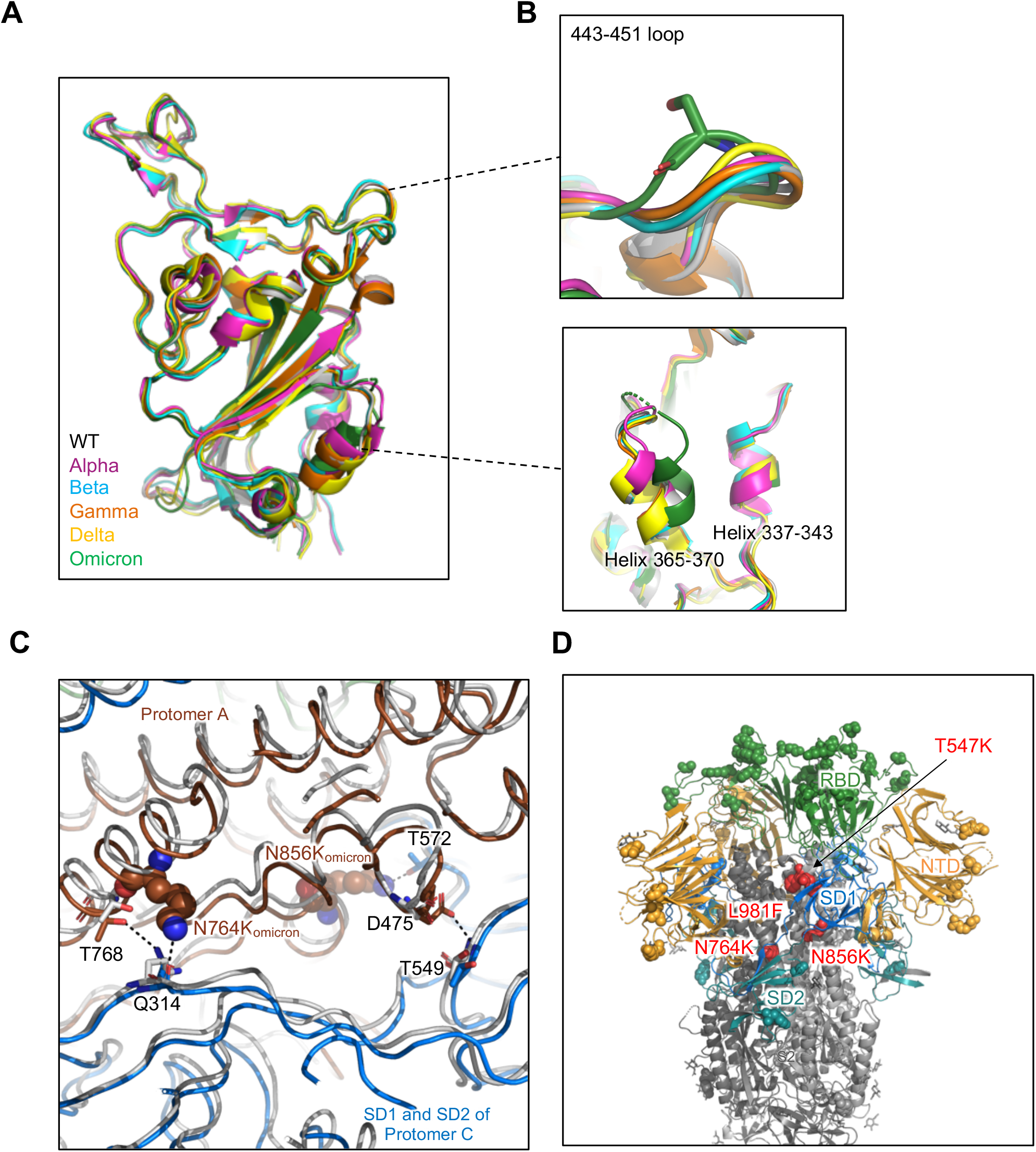
Structural comparison of SARS-CoV-2 omicron RBD with other VOCs, Related to Figure 3. (A) Superposition of RBDs of omicron and other VOCs. (B) Details of the conformational differences in omicron spike compared with WT and other VOCs. Up panel, 443-451 loop. Down panel, Helix 365-370 and Helix 337-443. (C) Ribbon diagram of omicron spike compared with WT. Black dashed line shows the hydrogen bonds. Residues mutated in omicron are shown as spheres, conserved residues in WT are shown as sticks. (D) Cartoon diagram of omicron spike. Mutations N764K, N856K, L981F, and T547K are labeled as red spheres.

**Figure S4.**
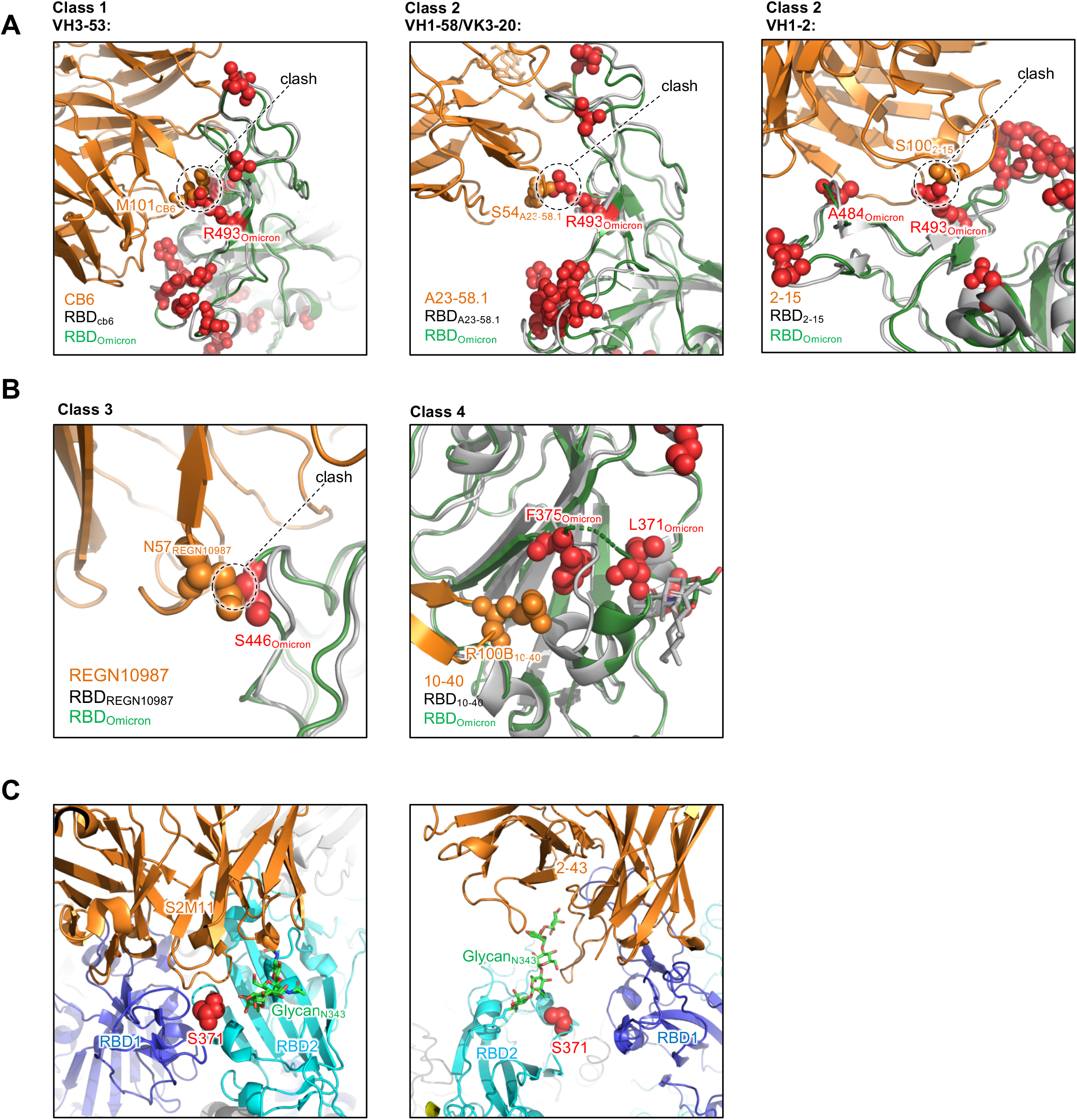
Escape mutations for different antibody classes in omicron spike, Related to Figure 4. (A) Cartoon diagram of the class 1 and class 2 antibody bound to omicron RBD. Left panel, VH3-53 derived antibody, CB6 as example. Middle panel, VH1-58/VK3-20 derived antibody, A23-58.1 as example. Right panel, VH1-2 derived antibody, 2-15 as example. Mutations in omicron RBD are shown as red spheres. Residues clashing with Omicron RBD are shown as orange spheres. (B) Cartoon diagram of the class 3 and class 4 antibodies bound to omicron RBD. REGN10987 and 10-40 as examples. (C) Cartoon diagram of 2 quaternary epitope recognizing antibodies that target N343 glycan. Left panel, S2M11. Right panel, 2-43.

**Table S1.**
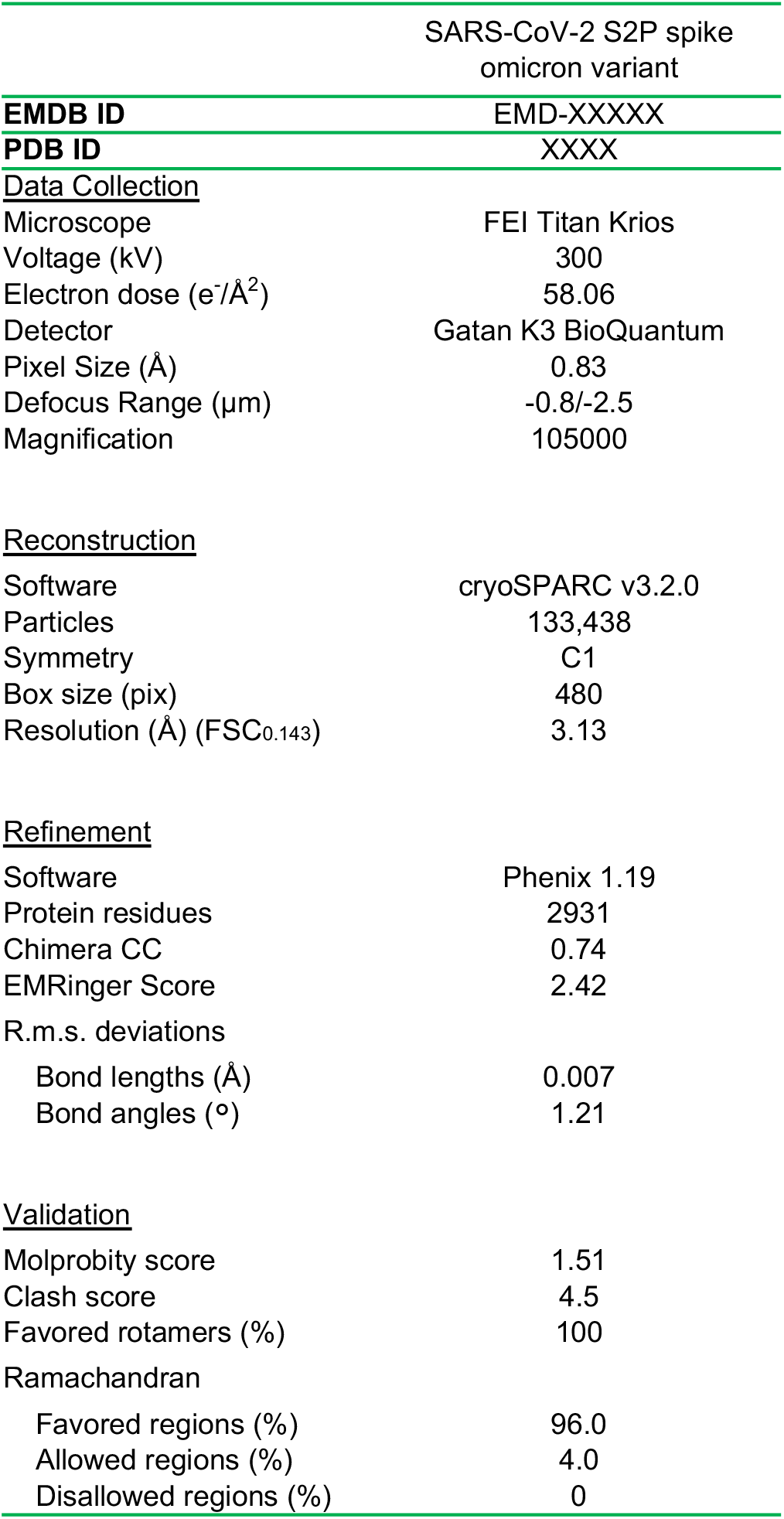
Cryo-EM Data Collection and Refinement Statistics. Related to Figure 1.

**Table S2.** Domain-Domain interface statistics for omicron spike compared with WT (PDB:7KRR). Related to Figure 2.

| Type | Domain | Interaction residues | Interface Area<br>(Å²) | ΔG<br>(kcal/mol) | Binding energy<br>(kcal/mol) | Hydrogen bonds | Salt bridges | Disulfide bonds |
| --- | --- | --- | --- | --- | --- | --- | --- | --- |
| WT | RBD:RBD | 6:8 | 136.7 | -0.1 | -0.1 | 0 | 0 | 0 |
| Omicron | RBD:RBD | 5:10 | 153.6 | -2.2 | -2.2 | 0 | 0 | 0 |
| WT | NTD:RBD | 10:9 | 193.8 | -0.4 | -0.4 | 0 | 0 | 0 |
| Omicron | NTD:RBD | 19:24 | 654.1 | -0.2 | -1.6 | 3 | 0 | 0 |
| WT | S2:S2 | 81:76 | 2778.1 | -30.4 | -42.9 | 23 | 6 | 0 |
| Omicron | S2:S2 | 89:76 | 2868.8 | -28.1 | -39.9 | 20 | 8 | 0 |
| WT | S2:SD1/SD2 | 50:48 | 1523.5 | -17.2 | -22.3 | 9 | 3 | 0 |
| Omicron | S2:SD1/SD2 | 48:49 | 1462.0 | -14.8 | -22 | 13 | 3 | 0 |

